# AnnotatorJ: an ImageJ plugin to ease hand-annotation of cellular compartments

**DOI:** 10.1101/2020.02.27.968362

**Authors:** Réka Hollandi, Ákos Diósdi, Gábor Hollandi, Nikita Moshkov, Péter Horváth

## Abstract

AnnotatorJ combines single-cell identification with deep learning and manual annotation. Cellular analysis quality depends on accurate and reliable detection and segmentation of cells so that the subsequent steps of analyses e.g. expression measurements may be carried out precisely and without bias. Deep learning has recently become a popular way of segmenting cells, performing unimaginably better than conventional methods. However, such deep learning applications may be trained on a large amount of annotated data to be able to match the highest expectations. High-quality annotations are unfortunately expensive as they require field experts to create them, and often cannot be shared outside the lab due to medical regulations.

We propose AnnotatorJ, an ImageJ plugin for the semi-automatic annotation of cells (or generally, objects of interest) on (not only) microscopy images in 2D that helps find the true contour of individual objects by applying U-Net-based pre-segmentation. The manual labour of hand-annotating cells can be significantly accelerated by using our tool. Thus, it enables users to create such datasets that could potentially increase the accuracy of state-of-the-art solutions, deep learning or otherwise, when used as training data.

## Introduction

Single-cell analysis pipelines begin with an accurate detection of the cells. Even though microscopy analysis software tools aim to become more and more robust to various experimental setups and imaging conditions, most lack efficiency in complex scenarios such as label-free samples or unforeseen imaging conditions (e.g. higher signal-to-noise ratio, novel microscopy or staining techniques), which opens up a new expectation of such software tools: adaptation ability (Hollandi et al., 2020). Another crucial requirement is to maintain ease of usage and limit the number of parameters the users need to fine-tune to match their exact data domain.

Recently, deep learning (DL) methods have proven themselves worthy of consideration in microscopy image analysis tools as they have also been successfully applied in a wider range of applications including but not limited to face detection (Taigman et al., 2014, Sun et al. 2014, Schroff et al. 2015), self-driving cars (Badrinarayanan et al., 2017, Redmon et al., 2016, Grigorescu et al., 2019) and speech recognition (Hinton et al., 2012). Caicedo et al. (Caicedo et al., 2019) and others (Hollandi et al 2020, Moshkov et al 2020) proved that single cell detection and segmentation accuracy can be significantly improved utilizing DL networks. The most popular and widely used deep convolutional neural networks (DCNNs) include Mask R-CNN (He et al., 2017): an object detection and instance segmentation network, YOLO (Redmon et al., 2016): a fast object detector and U-Net (Ronneberger et al., 2015): a fully convolutional network specifically intended for bioimage analysis purposes and mostly used for pixel classification. StarDist (Schmidt et al., 2018) is an instance segmentation DCNN optimal for convex or elliptical shapes (such as nuclei).

As robustly and accurately as they may perform, these networks rely on sufficient data, both in amount and quality, which tends to be the bottleneck of their applicability in certain cases such as single-cell detection. While in more industrial applications (see (Grigorescu et al., 2019) for an overview of autonomous driving) a large amount of training data can be collected relatively easily: see the cityscapes dataset (Cordts et al., 2016) (available at https://www.cityscapes-dataset.com/) of traffic video frames using a car and camera to record and potentially non-expert individuals to label the objects, clinical data is considerably more difficult, due to ethical constraints, and expensive to gather as expert annotation is required. Datasets available in the public domain such as BBBC (Ljosa et al., 2012) at https://data.broadinstitute.org/bbbc/, TNBC (Naylor et al., 2017, 2019) or TCGA (Cancer Genome Atlas Research Network, 2008; Kumar et al., 2017) and detection challenges including ISBI (Coelho et al., 2009), Kaggle (https://www.kaggle.com/ e.g. Data Science Bowl 2018, see at https://www.kaggle.com/c/data-science-bowl-2018), ImageNet (Russakovsky et al., 2015) etc. contribute to the development of genuinely useful DL methods; however, most of them lack heterogeneity of the covered domains and are limited in data size. Even combining them one could not possibly prepare their network/method to generalize well (enough) on unseen domains that vastly differ from the pool they covered. On the contrary, such an adaptation ability can be achieved if the target domain is represented in the training data, as proposed in (Hollandi et al 2020), where synthetic training examples are generated automatically in the target domain via image style transfer.

Eventually, similar DL approaches’ performance can only be increased over a certain level if we provide more training examples. The proposed software tool was created for this purpose: the expert can more quickly and easily create a new annotated dataset in their desired domain and feed the examples to DL methods with ease. The user-friendly functions included in the plugin help organize data and support annotation e.g. multiple annotation types, editing, classes etc. Additionally, a batch exporter is provided offering different export formats matching typical DL models’; supported annotation and export types are visualized on **Figure 1**, open source code is available at https://github.com/spreka/annotatorj under GNU GPLv3 license.

**Figure 1.**
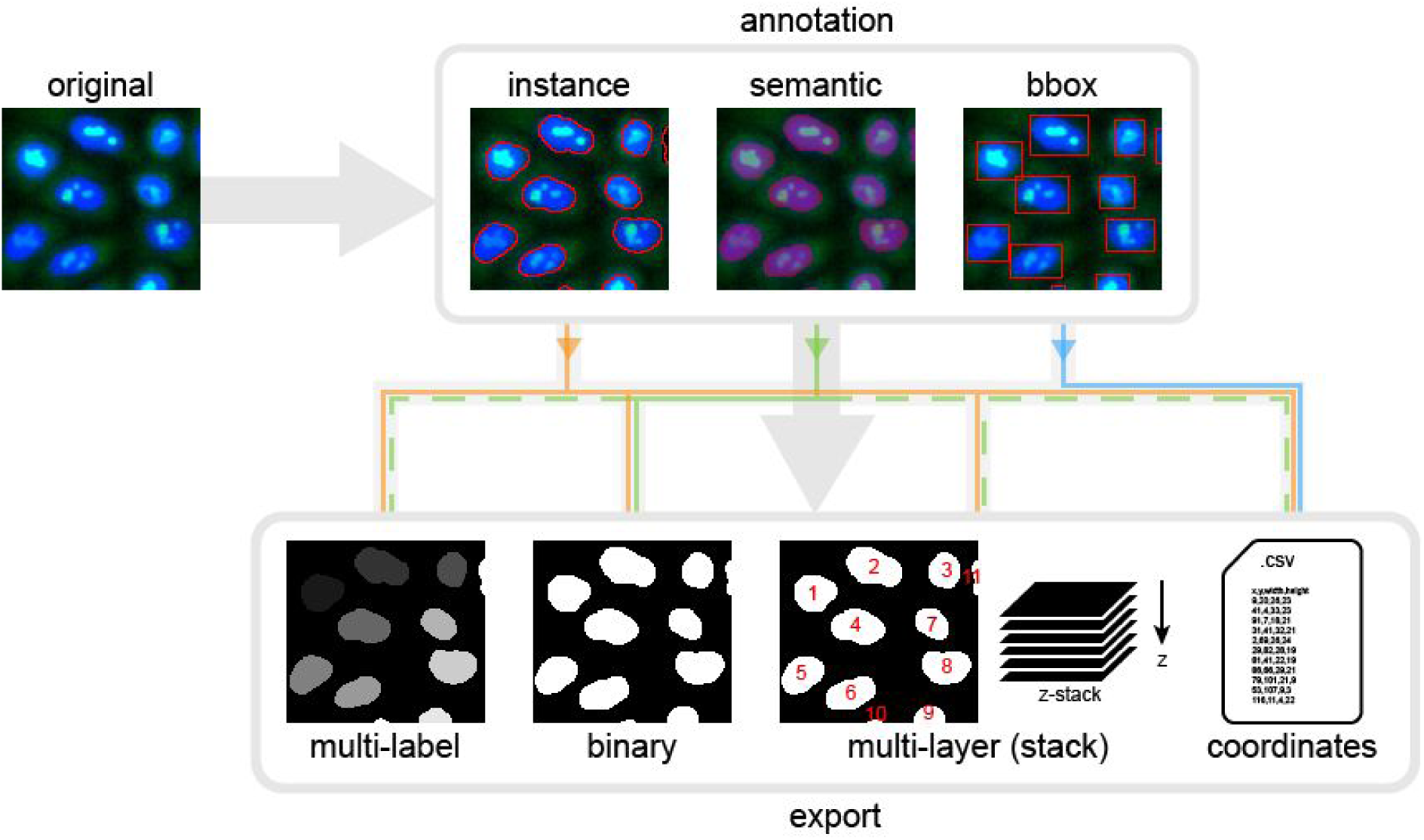
Annotation types. The top row displays our supported types of annotation: instance, semantic and bounding box (noted as ‘bbox’ on the figure) based on the same objects of interest, in this case nuclei, shown in red. Instances mark the object contours, semantic overlay shows the regions (area) covered, while bounding boxes are the smallest enclosing rectangles around the object borders. Export options are shown in the bottom row: multi-label, binary, multi-layer images and coordinates in a text file. Lines mark the supported export options for each annotation type by colours: orange for instance, green for semantic and blue for bounding box. Dashed lines indicate additional export options for semantic that should be used carefully.

We implemented the tool as an ImageJ (Abramoff et al 2004, Schneider et al 2012) plugin since ImageJ is frequently used by bioimage analysts, providing a familiar environment for users. While other software also provide means to support annotation e.g. by machine learning-based pixel-classification (see detailed comparison in Methods), AnnotatorJ is a lightweight, free, open source, cross-platform alternative. It can be easily installed via its ImageJ update site at https://sites.imagej.net/Spreka/ or run as a standalone ImageJ instance containing the plugin.

In AnnotatorJ we initialize annotations with DL pre-segmentation using U-Net to suggest contours from as little as a quickly drawn line over the object (see **Supplementary Material** and **Figure 2**). U-Net predicts pixels belonging to the target class with the highest probability within a small bounding box (a rectangle) around the initially drawn contour then a fine approximation of the true object boundary is calculated from connected pixels; this is referred to as the suggested contour. The user then manually refines the contour to create a pixel-perfect annotation of the object.

**Figure 2.**
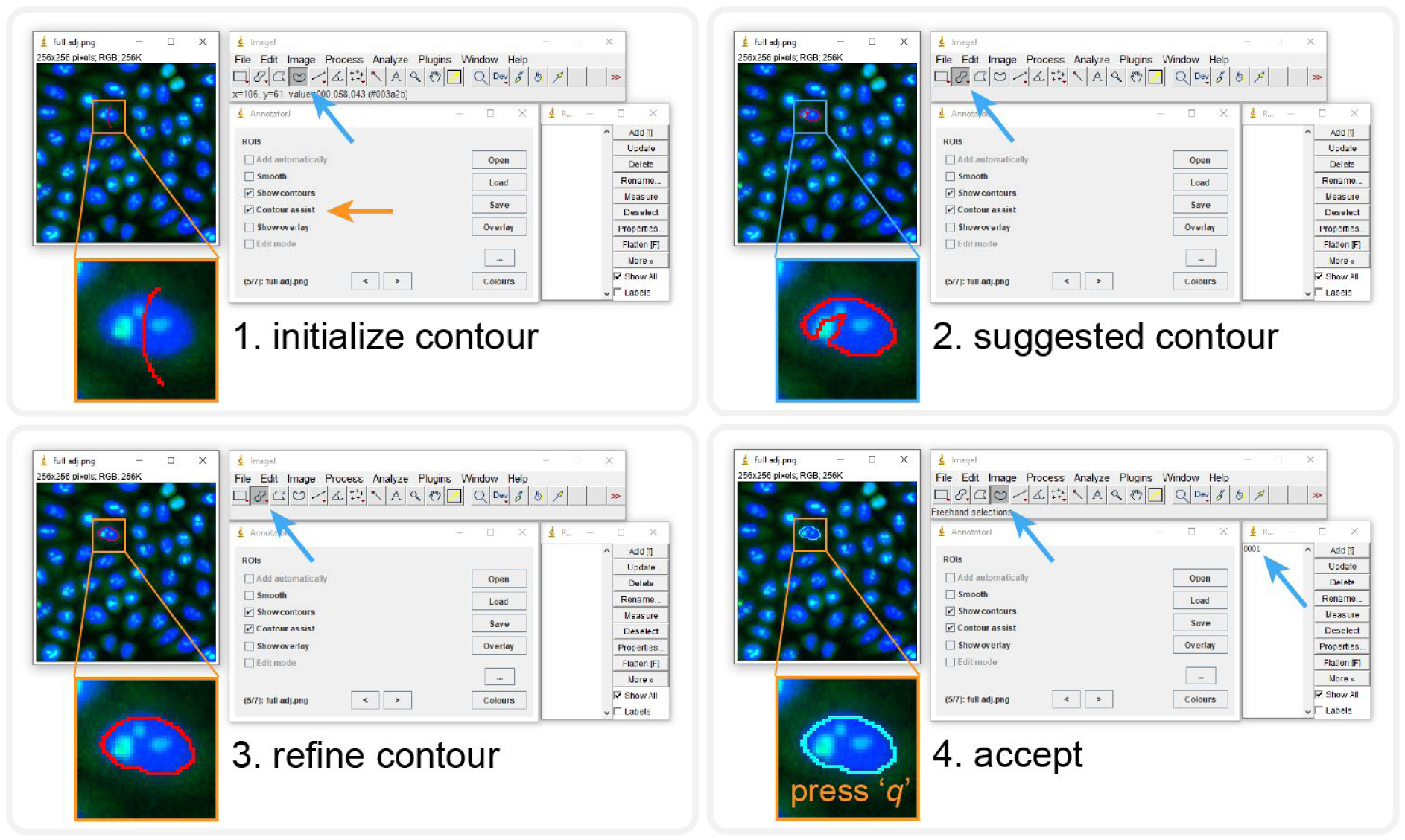
Contour assist mode of AnnotatorJ. The blocks show the order of steps, the given tool needed is automatically selected. User interactions are marked with orange arrows, automatic steps with blue. 1) Initialize the contour with a lazily drawn line, 2) the suggested contour appears (a window is shown until processing completes), brush selection tool is selected automatically, 3) refine the contour as needed, 4) accept it by pressing the key ‘q’ or reject with ‘Ctrl’+’delete’. Accepting adds the ROI to ROI Manager with a numbered label. See also **Supplementary Material** for a demo video (figure2.mov).

## Methods

### Motivation

We propose AnnotatorJ, an ImageJ (Abramoff et al 2004, Schneider et al 2012) plugin for the annotation and export of cellular compartments that can be used to boost DL models’ performance. The plugin is mainly intended for bioimage annotation but could possibly be used to annotate any type of objects on images (see **Fig.S2** for a general example). During development we kept in mind that the intended user should be able to get comfortable with the software really quickly and fasten the otherwise truly time-consuming and exhausting process of manually annotating single cells or their compartments (such as individual nucleoli, lipid droplets, nucleus or cytoplasm).

The performance of DL segmentation methods is significantly influenced by both the training data size and its quality. Should we feed automatically segmented objects to the network, errors present in the original data will be propagated through the network during training and bias the performance, hence such training data should always be avoided. Hand-annotated and curated data, however, will minimize the initial error boosting the expected performance increase on the target domain to which the annotated data belongs. NucleAIzer (Hollandi et al., 2020) showed increase in nucleus segmentation accuracy when a DL model was trained on synthetic images generated from ground truth annotations instead of pre-segmented masks.

### Features

AnnotatorJ helps organize the input and output files by automatically creating folders and matching file names to the selected type and class of annotation. Currently, the supported annotation types are 1) instance, 2) semantic and 3) bounding box; see **Figure 1**. Each of these are typical inputs of DL networks; instance annotation provides individual objects separated by their boundaries (useful in case of e.g. clumped cells of cancerous regions) and can be used to provide training data for instance segmentation networks such as Mask R-CNN (He et al., 2017). Semantic annotation means foreground-background separation of the image without distinguishing individual objects (foreground), a typical architecture using such segmentations is U-Net (Ronneberger et al., 2015). And finally, bounding box annotation is done by identifying the object’s bounding rectangle, and is generally used in object detection networks (like YOLO (Redmon et al., 2016) or R-CNN (Girshick et al., 2014)).

Semantic annotation is done by painting areas on the image overlay. All necessary tools to operate a given function of the plugin are selected automatically. Contour or overlay colours can be selected from the plugin window. For a detailed description and user guide please see the documentation of the tool (available at https://github.com/spreka/annotatorj repository).

Annotations can be saved to default or user-defined “classes” corresponding to biological phenotypes (e.g. normal or cancerous) or object classes – used as in DL terminology (such as person, chair, bicycle etc.), and later exported in a batch by class. Phenotypic differentiation of objects can be supported by loading a previously annotated class’s objects for comparison as overlay to the image and toggling their appearance by a checkbox.

We use the default ImageJ ROI Manager to handle instance annotations as individual objects (ROI(s): region of interest). Annotated objects can be added to the ROI list automatically (without the bound keystroke “t” as defined by ROI Manager) when the user releases the mouse button used to draw the contour by checking its option in the main window of the plugin. This ensures that no annotation drawn is missing from the ROI list.

Contour editing is also possible in our plugin using “Edit mode” (by selecting its checkbox) in which the user can select any already annotated object on the image by clicking on it, then proceed to edit the contour and either apply modifications with the shortcut ‘Ctrl’+’q’, discard them with ‘escape’ or delete the contour with ‘Ctrl’+’delete’. The given object selected for edit is highlighted in inverse contour colour.

Object-based classification is also possible in “Class mode” (via its checkbox): similarly to “Edit mode”, ROIs can be assigned to a class by clicking on them on the image which will also update the selected ROI’s contour to the current class’s colour. New classes can be added and removed, their class colour can be changed. A default class can be set for all unassigned objects on the image. Upon export (using either the quick export button “[^]” in the main window or the exporter plugin) masks are saved by classes.

In the options (button “…” in the main window) the user can select to use either U-Net or a classical region-growing method to initialize the contour around the object marked. Currently only instance annotation can be assisted.

### Contour suggestion using U-Net

Our annotation helper feature “*Contour assist*” (see **Figure 2**) allows the user to work on initialized object boundaries by roughly marking an object’s location on the image which is converted to a well-defined object contour via weighted thresholding after a U-Net (Ronneberger et al., 2015) model trained on nucleus or other compartment data predicts the region covered by the object. We refer this as the *suggested contour* and expect the user to refine the boundaries to match the object border precisely. The suggested contour can be further optimized by applying *active contour* (AC) (Kass et al., 1988) to it. We aim to avoid fully automatic annotation (as previously argued) by only enabling one object suggestion at a time and requiring manual interaction to either refine, accept or reject the suggested contour. These operations are bound to keyboard shortcuts for convenience (see **Fig.2**). When using the *Contour assist* function automatic adding of objects is not available to encourage the user to manually validate and correct the suggested contour as needed.

On **Figure 2** we demonstrate *Contour assist* using a U-Net model trained on versatile microscopy images of nuclei in Keras and on a fluorescent microscopy image of a cell culture where the target objects, nuclei are labelled with DAPI (in blue). This model is provided at https://github.com/spreka/annotatorj/releases/tag/v0.0.2-model in the open-source code repository of the plugin.

Contour suggestions can be efficiently used for proper initialization of object annotation, saving valuable time for the expert annotator by suggesting a nearly perfect object contour that only needs refinement (as shown on **Figure 2**). Using a U-Net model accurate enough for the target object class the expert can focus on those image regions where the model is rather uncertain (e.g. around the edges of an object or the separating line between adjacent objects) and fine-tune the contour accurately while sparing considerable effort on more obvious regions (like an isolated object on simple background) by accepting the suggested contour after marginal correction.

The framework of the chosen U-Net implementation, Deeplearning4j (DL4J, available at http://deeplearning4j.org/ or https://github.com/eclipse/deeplearning4j), supports Keras model import, hence custom, application-specific models can be loaded in the plugin easily by either training them in DL4J (Java) or Python (Keras) and saving the trained weights and model configuration in .h5 and .json files. This vastly extends the possible fields of application for the plugin to general object detection or segmentation tasks (see **Supplementary Material** and **Figures S2-S3**).

### Exporter

The annotation tool is supplemented by an exporter, AnnotatorJExporter plugin also available in the package. It was optimized for the batch export of annotations created by our annotation tool. For consistency, one class of objects can be exported at a time. We offer 4 export options: 1) multi-labelled, 2) multi-layered and 3) semantic images and 4) coordinates; see **Figure 1**. Instance annotations are typically expected to be exported as multi-labelled (instance-aware) or multi-layered (stack) grayscale images, the latter of which is useful for handling overlapping objects such as cytoplasms in cell culture images. Semantic images are binary foreground-background images of the target objects while coordinates (top-left corner (x,y) of the bounding rectangle appended by its width and height in pixels) can be useful training data for object detection applications including astrocyte localization (Suleymanova et al., 2018) or in a broader aspect, face detection (Taigman et al., 2014). All export options are supported for semantic annotation, however, we note that in instance-aware options (multi-labelled or multi-layered mask and coordinates) only such objects are distinguished whose contours do not touch on the annotation image.

### OpSeF compatibility

OpSeF (Open Segmentation Framework, Rasse et al., 2020) is an interactive python notebook-based framework (available at https://github.com/trasse/OpSeF-IV) that allows users to easily try different DL segmentation methods in customizable pipelines. We extended AnnotatorJ to support the data structure and format used in OpSeF to allow seamless integration in these pipelines, so users can manually modify, create or classify objects found by OpSeF in AnnotatorJ, then export the results in a compatible format for further use in the former software. User guide is provided in the documentation of https://github.com/trasse/OpSeF-IV.

## Results

### Performance evaluation

We quantitatively evaluated annotation performance and speed in AnnotatorJ (see **Figures 3-4**) with the help of three annotators who had experience in cellular compartment annotation. Both annotation accuracy and time was measured on the same two test sets: a nucleus and a cytoplasm image set (see also **Figure S1** and **Supplementary Material**). Both test sets contained images of various experimental conditions, including fluorescently labelled- and brightfield-stained samples, tissue section and cell culture images. We compared the effectiveness of our plugin using *Contour assist* mode to only allowing the use of *Automatic adding*. Even though the latter is also a functionality of AnnotatorJ, it ensured that the measured annotation times correspond to a single object each. Without this option the user must press the key “t” after every contour drawn to add it to the ROI list which can be unintendedly missed, increasing its time as the same contour must be drawn again.

**Figure 3.**
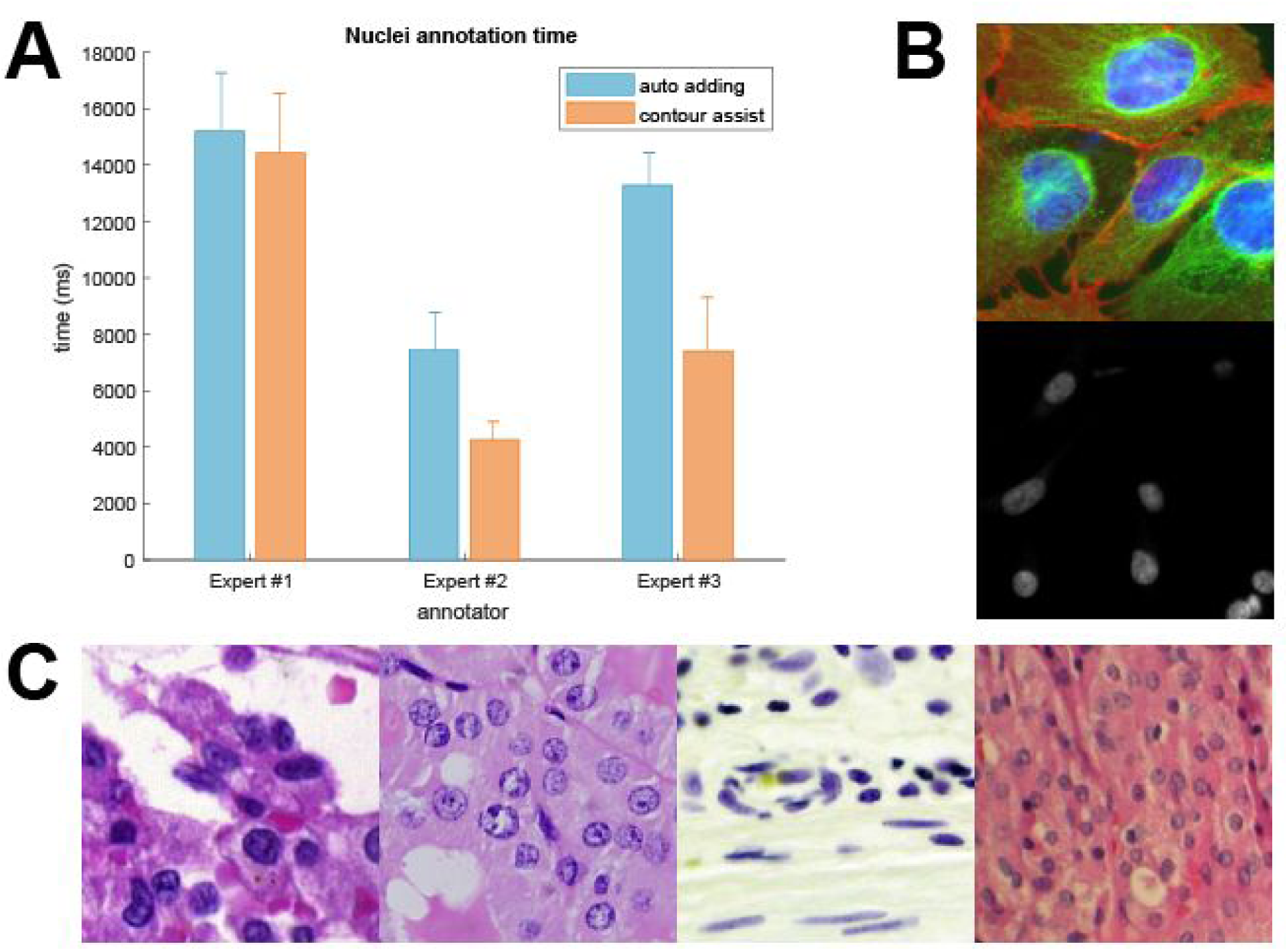
Annotation times on nucleus images. AnnotatorJ was tested on sample microscopy images (both fluorescent and brightfield, as well as cell culture and tissue section images), annotation time was measured on a per-object (nucleus) level. Bars represent the mean annotation times on the test image set, error bars show SEM (standard error of the mean). Orange corresponds to Contour assist mode and blue to only allowing the Automatic adding option. A) Nucleus test set annotation times, B) example cell culture test images, C) example histopathology images. Images shown on B-C) are 256×256 crops of original images. Some images are courtesy of Kerstin Elisabeth Dörner, Andreas Mund, Viktor Honti and Hella Bolck.

**Figure 4.**
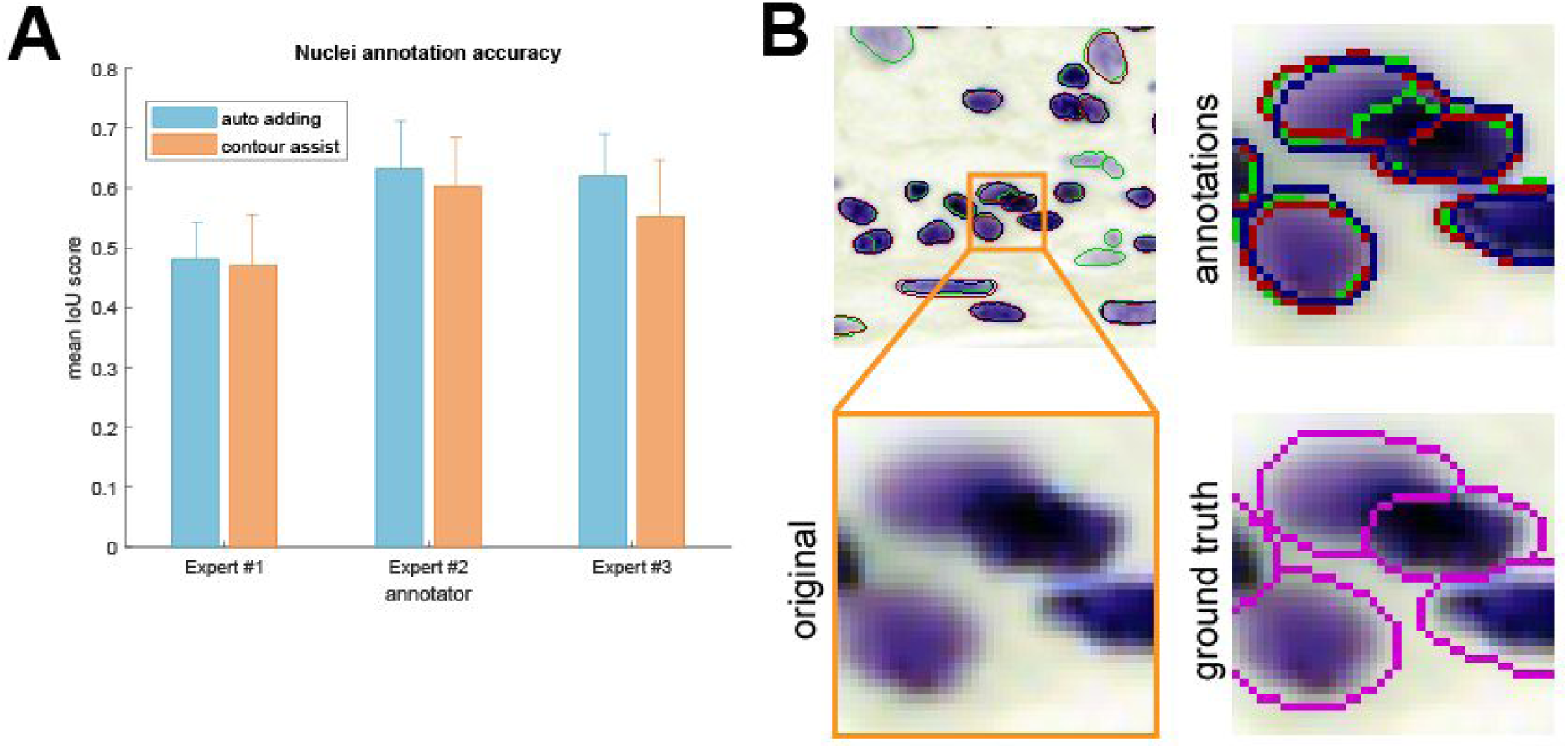
Annotation accuracies. Annotations created in the same test as the times measured in Figure 3 were evaluated using mean IoU scores for the nucleus test set. Error bars show SEM, colours are as in Figure 3. A) Nucleus test set accuracies, B) example contours drawn by our expert annotators. Inset highlighted in orange is zoomed in showing the original image, annotations marked in red, green and blue corresponding to experts #1-3, respectively, and ground truth contours (in magenta) are overlayed on the original image for comparison.

For the annotation time test presented on **Figure 3** we measured the time passed between adding new objects to the annotated object set in ROI Manager for each object, then averaged the times for each image and each annotator, respectively. Time was measured in the Java implementation of the plugin in milliseconds. **Figures 3-4** show SEM (standard error of the mean) error bars for each mean measurement (see **Supplementary Material** for details).

In the case of annotating cell nuclei, results confirm that hand-annotation tasks can be significantly accelerated using our tool. Each three annotators were faster by using *Contour assist*, two of them nearly double their speed.

To ensure efficient usage of our plugin in annotation assistance, we also evaluated the accuracies achieved in each test case by calculating mean intersection over union (IoU) scores of the annotations as segmentations compared to ground truth masks previously created by different expert annotators. We used the mean IoU score defined in the Data Science Bowl 2018 competition (https://www.kaggle.com/c/data-science-bowl-2018/overview/evaluation) and in (Hollandi et al 2020):

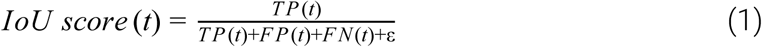

IoU determines the overlapping pixels of the segmented mask with the ground truth mask (intersection) compared to their union. IoU score is calculated at 10 different thresholds from 0.5 to 0.95 with 0.05 steps, at each threshold true positive (TP), false positive (FP) and false negative (FN) objects are counted. An object is considered true positive if its IoU is greater than the given threshold *t*. IoU scores calculated at all 10 thresholds were finally averaged to yield a single IoU score for a given image in the test set.

An arbitrarily small ε = 10^-40^ value was added to the denominators for numerical stability. (1) is a modified version of mean average precision (mAP) typically used to describe the accuracy of instance segmentation approaches. Precision is formulated as

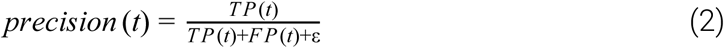

Nucleus and cytoplasm image segmentation accuracies were averaged over the test sets, respectively. We compared our annotators using and not using *Contour assist* mode (**Figure 4**). The results show greater inter-expert than intra-expert differences allowing us to conclude that the annotations created in AnnototarJ are nearly as accurate as free-hand annotations.

### Export evaluation

As the training data annotation process for deep learning applications requires the annotated objects to be exported in a manner that DL models can load them, which typically covers the four types of export options offered in AnnotatorJExporter, it is also important to investigate the efficiency of export. We measured export times similarly to annotation times. For the baseline results each object defined by their ROIs was copied to a new empty image, filled and saved to create a segmentation mask image file. Exportation from AnnotatorJExporter was significantly faster and only required a few clicks: it took 4 orders of magnitude less time to export the annotations (about 60 ms). Export times reported correspond to a randomly selected expert so that computer hardware specifications remain the same.

### Comparison to other tools and software packages

The desire to collect annotated datasets has arisen with the growing popularity and availability of application specific DL methods. Object classification on natural images (photos) and face recognition are frequently used examples of such applications in computer vision. We discuss some of the available software tools created for image annotation tasks and compare their feature scope in the following table **(Table 1, see also Table S1)** and in **Supplementary Material**.

**Table 1.**
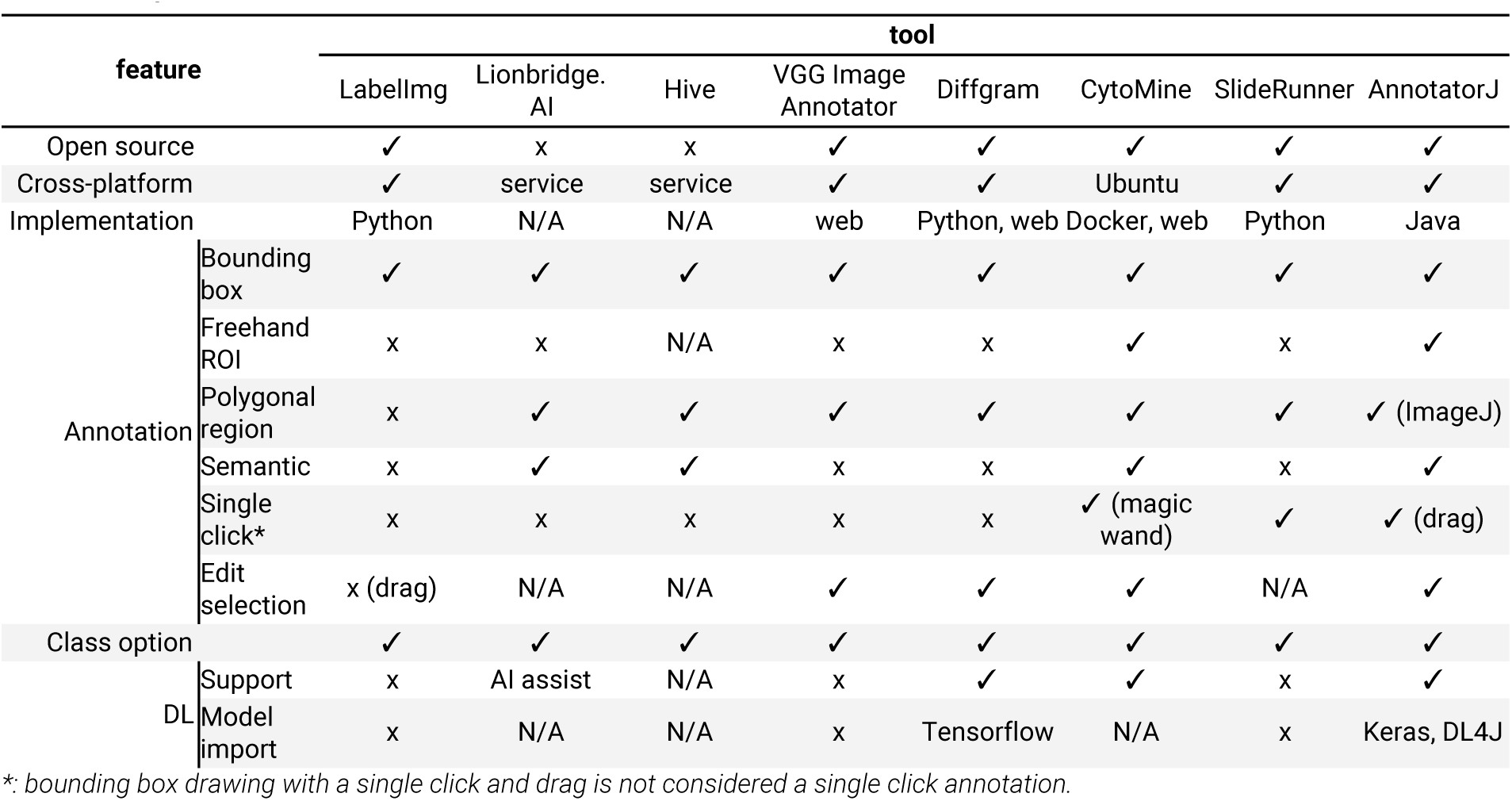
Comparison of annotation software tools.

We collected our list of methods to compare following (Morikawa 2019) and (“The best image annotation platforms”, 2018). While there certainly is a considerable amount of annotation tools for object detection purposes, most of them are not open source. We included Lionbridge.AI (https://lionbridge.ai/services/image-annotation/) and Hive (https://thehive.ai/), two service-based solutions because of their wide functionality and artificial intelligence support. Both of them work in a project-management way and outsource the annotation task to enable fast and accurate results. Their main application spectra covers more general object detection tasks like classification of traffic video frames. LabelImg (https://github.com/tzutalin/labelImg) on the other hand, as well as the following tools, is open source but offers a narrower range of annotation options and lacks machine learning support making it a lightweight but free alternative. VGG Image Annotator (Dutta and Zisserman, 2019) comes on a web-based platform therefore makes it really easy for the user to familiarize with the software. It enables multiple types of annotation with class definition. Diffgram (https://diffgram.com/) is available both online and as a locally installable version (Python) and adds DL support which speeds up the annotation process significantly - that is, provided the intended object classes are already trained and the DL predictions only need minor edit. A similar, also web-based approach is provided by supervise.ly (https://supervise.ly/, see in **Supplementary Material**) which is free for research purposes. Even though web-hosted services offer a convenient solution for training new models (if supported), handling sensitive clinical data may be problematic. Hence, locally installable software are more desirable in biological and medical applications. A software closer to the bioimage analyst community is CytoMine (Marée et al., 2016; Rubens et al., 2019), a more general image processing tool with a lot of annotation options that also provides DL support and has a web interface. SlideRunner (Aubreville et al., 2018) was created for large tissue section (slide) annotation specifically but similar to others it does not integrate machine learning methods to help annotation and rather focuses on the classification task.

AnnotatorJ on the other hand, as an ImageJ (Fiji) plugin should provide a familiar environment for bioimage annotators to work in. It offers all the functionality available in similar tools (such as different annotation options: bounding box, polygon, free hand drawing, semantic segmentation and editing them) while also incorporates support for a popular DL model, U-Net. Furthermore, any user-trained Keras model can be loaded into the plugin with ease because of the DL4J framework, extending its use cases to general object annotation tasks (see **Fig. S2** and **Supplementary Material**). Due to its open source implementation the users can modify or extend the plugin to even better fit their needs. Additionally, as an ImageJ plugin it requires no software installation, can be downloaded inside ImageJ/Fiji (via its update site, https://sites.imagej.net/Spreka/) or run as a standalone ImageJ instance with this plugin.

We also briefly discuss ilastik (Sommer et al., 2011; Berg et al., 2019) and Suite2p (Pachitariu et al., 2017) in the **Supplementary Material** since they are not primarily intended for annotation purposes. Two ImageJ plugins that offer manual annotation and machine learning-generated outputs, Trainable Weka Segmentation (Arganda-Carreras et al., 2017) and LabKit (Arzt 2017) are also detailed in the **Supplementary Material**.

## Discussion

We presented an ImageJ plugin, AnnotatorJ for convenient and fast annotation and labelling of objects on digital images. Multiple export options are also offered in the plugin.

We tested the efficiency of our plugin with three experts on two test sets comprising of nucleus and cytoplasm images. We found that our plugin accelerates the hand-annotation process on average and offers up to four orders of magnitude faster export. By integrating the DL4J Java framework for U-Net contour suggestion in *Contour assist* mode any class of object can be annotated easily: the users can load their own custom models for the target class.

## Materials and methods

### ImageJ

ImageJ (or Fiji: Fiji is just ImageJ (Schindelin et al., 2012)) is an open-source, cross-platform image analysis software tool in Java that has been successfully applied in numerous bioimage analysis tasks (segmentation (Arganda-Carreras et al., 2017; Legland et al., 2016), particle analysis (Abramoff et al. 2004) etc.) and is supported by a broad range of community, comprising of biomage analyst end users and developers as well. It provides a convenient framework for new developers to create their custom plugins and share them with the community. Many typical image analysis pipelines have already been implemented as a plugin, e.g. U-Net segmentation plugin (Falk et al., 2019) or StarDist segmentation plugin (Schmidt et al., 2018).

### U-Net implementation

We used the Deeplearning4j (DL4J, http://deeplearning4j.org/) implementation of U-Net in Java. DL4J enables building and training custom DL networks, preparing input data for efficient handling and supports both GPU and CPU computation throughout its ND4J library.

The architecture of U-Net was first developed by Ronneberger *et al* (Ronneberger et al., 2015) and was designed to learn medical image segmentation on a small training set when limited amount of labelled data is available, which is often the case in biological contexts. To handle touching objects as often the case in nuclei segmentation, it uses a weighted cross entropy loss function to enhance the object-separating background pixels.

### Region-growing

A classical image processing algorithm, region-growing (Adams and Bischof, 1994; Haralick and Shapiro, 1985) starts from initial seed points or objects and expands the regions towards the object boundaries based on the intensity changes on the image and constraints on distance or shape. We used our own implementation of this algorithm.

## Supporting information

Contour assist mode of AnnotatorJ. Related to Figure 2.

## Abbreviations

DL: deep learning
ROI: region of interest
DL4J: Deeplearning4j
IoU: intersection over union

## Acknowledgements

R.H., N.M., A.D., and P.H. acknowledge support from the LENDULET-BIOMAG Grant (2018-342), from the European Regional Development Funds (GINOP-2.3.2-15-2016-00006, GINOP-2.3.2-15-2016-00026, GINOP-2.3.2-15-2016-00037), from the H2020-discovAIR (874656), and from Chan Zuckerberg Initiative, Seed Networks for the HCA-DVP. We thank Krisztián Koós for the technical help, and Máté Görbe and Tamás Monostori for testing the plugin.

## Supplementary Material

### Test data

We tested the efficiency in terms of both annotation time and accuracy on two test image sets: nucleus and cytoplasm. A sample of the test sets is displayed on **Figures 3B-C** and **S1B-C**; images are collected from publicly available datasets and our collaborators (see sources in (Hollandi et al. 2020)). We collected images from various modalities including different microscopy imaging (brightfield, fluorescent), sample origin (cell lines, histological tissue sections), staining technique (label-free, immunohistochemical or fluorescent markers for specific cellular compartments) that show the target object (nucleus or cytoplasm) in a heterogeneous environment so that we can test our plugin in general cases.

### Evaluation

We used SEM (standard error of the mean) in our evaluation of annotation time and accuracy which is formulated as

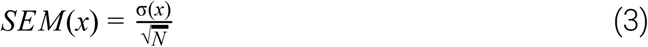

where the sample is denoted as *x*, σ(*x*) marks the standard deviation of *x* and *N* is the number of elements in *x*.

We also tested annotation on a cytoplasm test image set containing more complex regions e.g. overlapping objects (see **Figure S1**). Our cytoplasm model aided annotation, however, due to its semantic nature it could not always separate touching objects properly hence the increased annotation time in certain cases.

**Supplementary Figure S1.**
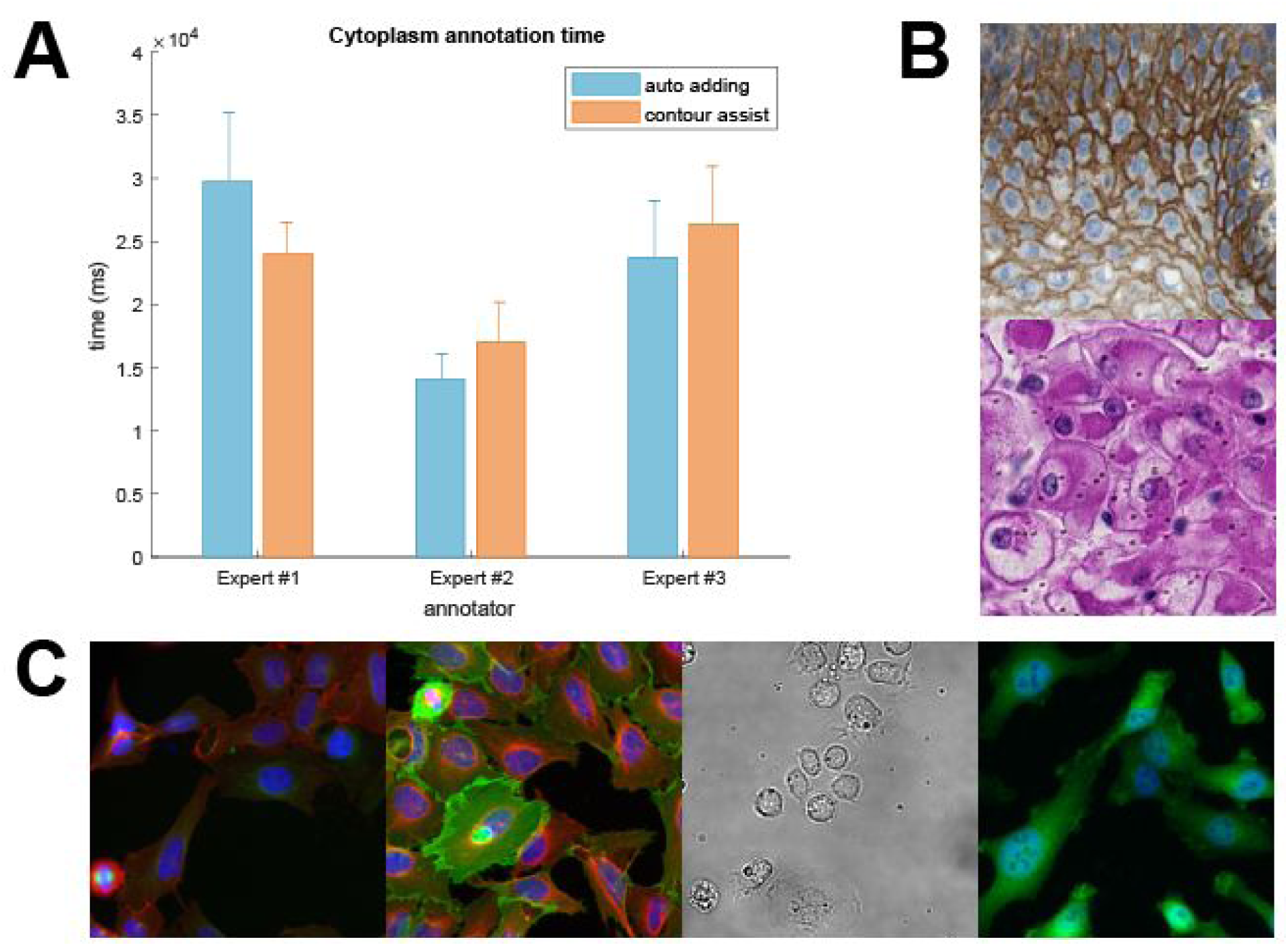
Annotation times on cytoplasm images. AnnotatorJ was tested on sample microscopy images (both fluorescent and brightfield, as well as cell culture and tissue section images), annotation time was measured on a per-object (cytoplasm) level. Bars represent the mean annotation times on the test image set, error bars show SEM (standard error of the mean). Orange corresponds to Contour assist mode and blue to only allowing the Automatic adding option. A) Cytoplasm annotation times, B) example histopathology test images, C) example cell culture images. Images shown on B-C) are 256×256 crops of original images. Some images are courtesy of Kerstin Elisabeth Dörner, Andreas Mund, Viktor Honti and Hella Bolck.

As any user-defined U-Net model can be easily imported in AnnotatorJ via DL4J, we tested general applicability on the Cityscapes dataset (Cordts et al., 2016): we trained a custom model on the car object class. Expert annotators were not needed for testing on these everyday objects. **Figure S2** shows annotation times and example images. Acceleration in annotation time is also significant (similarly to the nucleus test set); contour suggestion helps the most in complex-shaped regions such as outside mirrors and bumpers.

**Supplementary Figure S2.**
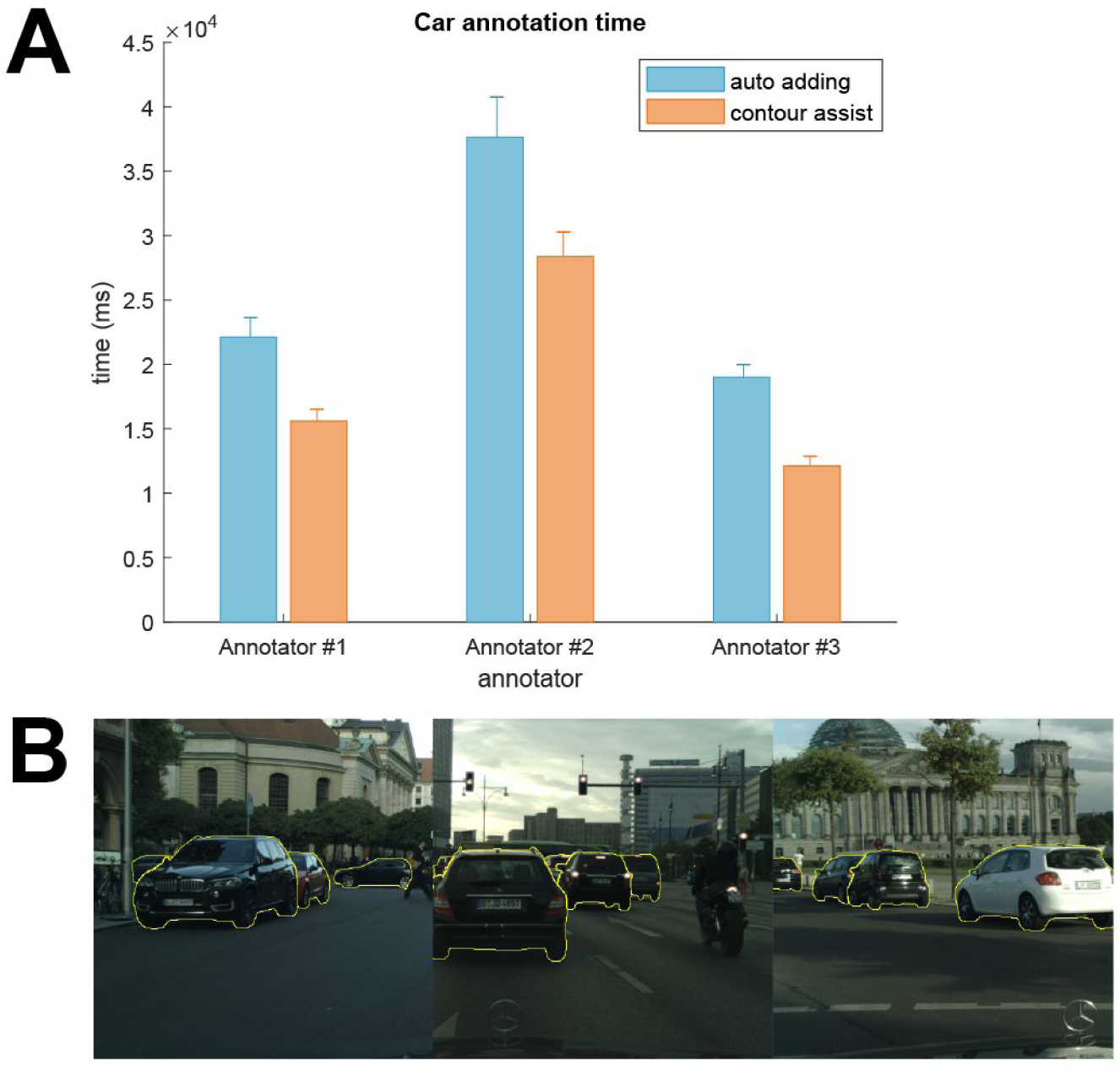
Annotation times on car images. AnnotatorJ was tested on sample images of the Cityscapes dataset (Cordts et al., 2016), annotation time was measured on a per-object (car) level. Bars represent the mean annotation times on the test image set, error bars show SEM (standard error of the mean). Orange corresponds to Contour assist mode and blue to only allowing the Automatic adding option. A) Car annotation times, B) example images as 1024×1024 crops of original images outlined by a randomly selected annotator.

Additionally, we tested AnnotatorJ using a model trained on electron microscopy (EM) data from the ISBI 2012 challenge (Arganda-Carreras et al., 2015) (fetched from https://github.com/zhixuhao/unet, which repository was also used to train our U-Net models in Keras) to help annotate neuronal cells (see **Fig. S3**). The complex shapes these structures had also emphasized the efficiency of our contour suggestion.

**Supplementary Figure S3.**
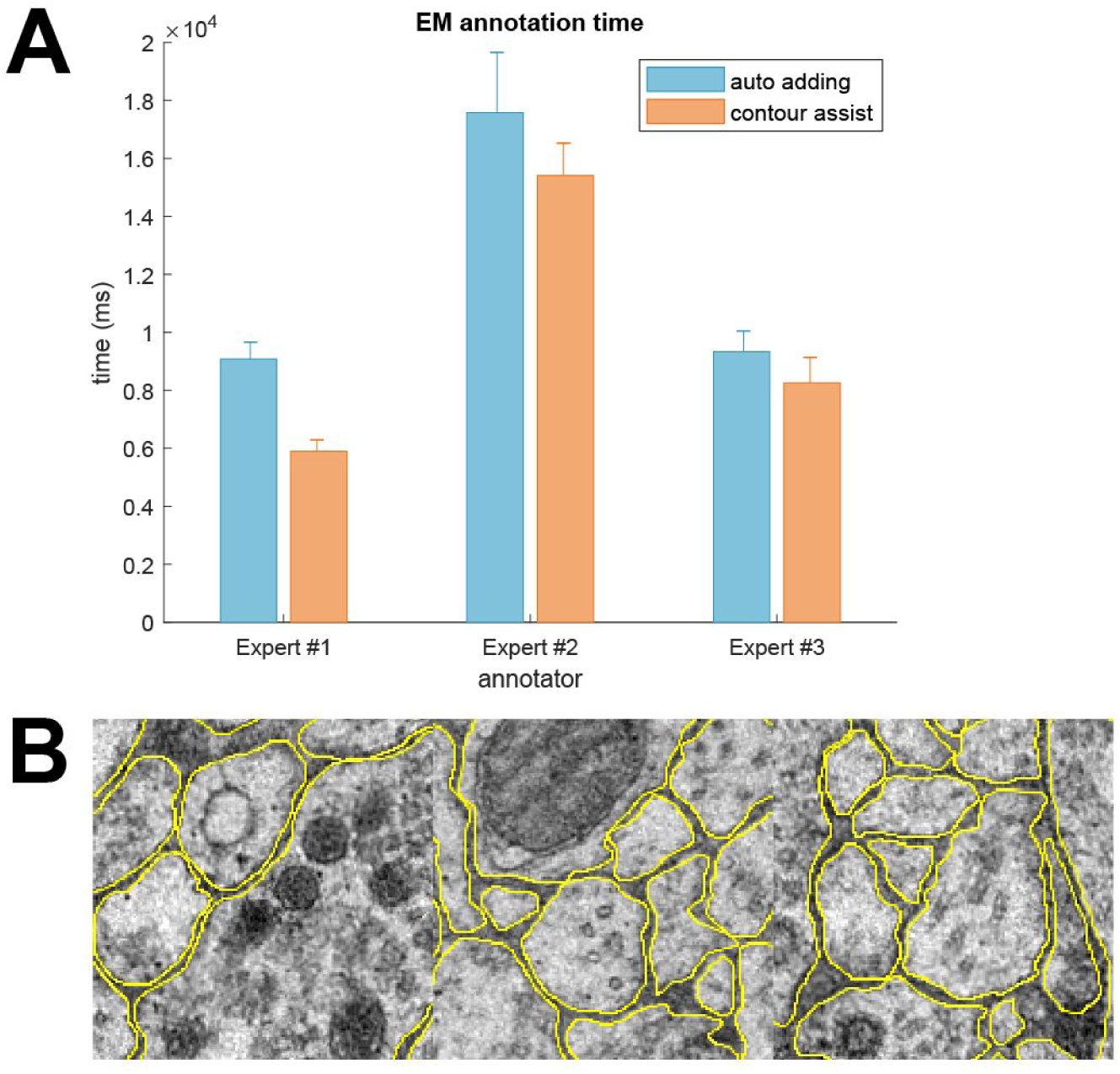
Annotation times on electron microscopy (EM) images. AnnotatorJ was tested on sample images of the ISBI 2012 challenge (Arganda-Carreras et al., 2015), annotation time was measured on a per-object (cell) level. Bars represent the mean annotation times on the test image set, error bars show SEM (standard error of the mean). Orange corresponds to Contour assist mode and blue to only allowing the Automatic adding option. A) EM annotation times, B) example images as 256×256 crops of original images outlined by a randomly selected expert.

## Additional methods

### Active contours (AC)

Active contours (Kass et al., 1988) is a generally well applicable image processing technique to fit an initial contour to the boundaries of an object by evolving a so-called *snake* contour driven by 2 energy functions as follows.

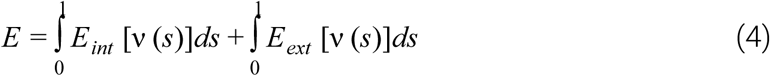

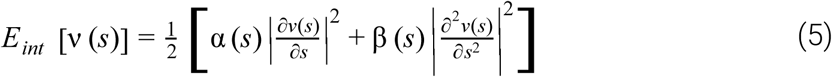

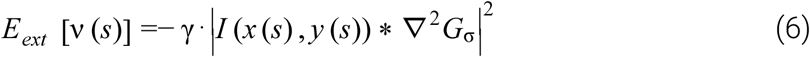

(4) is the combination of the internal (5) and external (6) energy functions, and is minimized during the iterative evolution process. The internal energy is responsible for extending and smoothing the contour while the external energy restrictively drives it towards the edges of the image which are enhanced by the LoG (Laplacian of Gaussian) filter-convolved image (data term). α, β and γ are the regularization parameters.

### Gradient vector flow (GVF)

GVF was first published in (Xu and Prince 1997) as a method to define a vector field based on the image gradient, that is the derivatives of the image, as enhanced by edges. It is attracted to bright regions (ideally edges) on the image.

## Additional annotation tools

Since open source or local software is of primary interest in medical research (see in Introduction and *Comparison to other tools and software packages*), we extend our previous list of compared tools here.

We briefly discussed supervise.ly (https://supervise.ly/) above, a service-based web-hosted solution for data labelling with DL model support. Training of various DL models including Mask R-CNN, U-Net, YOLO and others is possible. It supports annotation as bounding box, semantic- and polygonal drawing, editing points in polygons and class labelling. However, its free community edition intended for research purposes provides limited functionality.

FastAnnotationTool (https://github.com/christopher5106/FastAnnotationTool) is a lightweight, open source bounding box annotation tool with editing option in C++.

Make Sense (https://www.makesense.ai/) is another open source labelling tool that also adds DL support for annotation suggestion (using pre-trained models) and labelling. Point, polygon and bounding box annotation is possible, and classes can be assigned. Implemented in TypeScript, it offers both a web-based and a local (https://github.com/SkalskiP/make-sense) version.

ilastik (Sommer et al., 2011; Berg et al., 2019) is a pixel-based classification software that can be trained via brush strokes on the image. It can export semantic segmentation masks and their probability maps as well. However, these are returned by machine learning algorithms; users can annotate regions (pixel-based) with a brush or objects (instances) by single clicks to train them.

Suite2p (Pachitariu et al., 2017) was designed for a specific task: neuronal two-photon microscopy image analysis. It has a pipeline that can be modified in each step so that it fits the user’s data, automatic detection and classification of ROIs (cells) can be manually curated.

LabKit (Arzt 2017) is an ImageJ plugin for machine learning-supported labelling that similarly to the Trainable Weka Segmentation (ImageJ) plugin (Arganda-Carreras et al., 2017) and ilastik (Sommer et al., 2011; Berg et al., 2019) uses user-defined regions marked by brush strokes to train a machine learning algorithm and automatically segment image pixels to the defined classes. Both plugins offer several machine learning algorithms implemented in the Weka framework (Frank et al., 2016). Trainable Weka Segmentation can save semantic segmentation masks or their probability maps (like ilastik) while LabKit can also save manually drawn objects as multi-labelled .tiff files (like AnnotatorJExporter).

We collected the sources of annotation tools in Table S1.

**Table S1.**
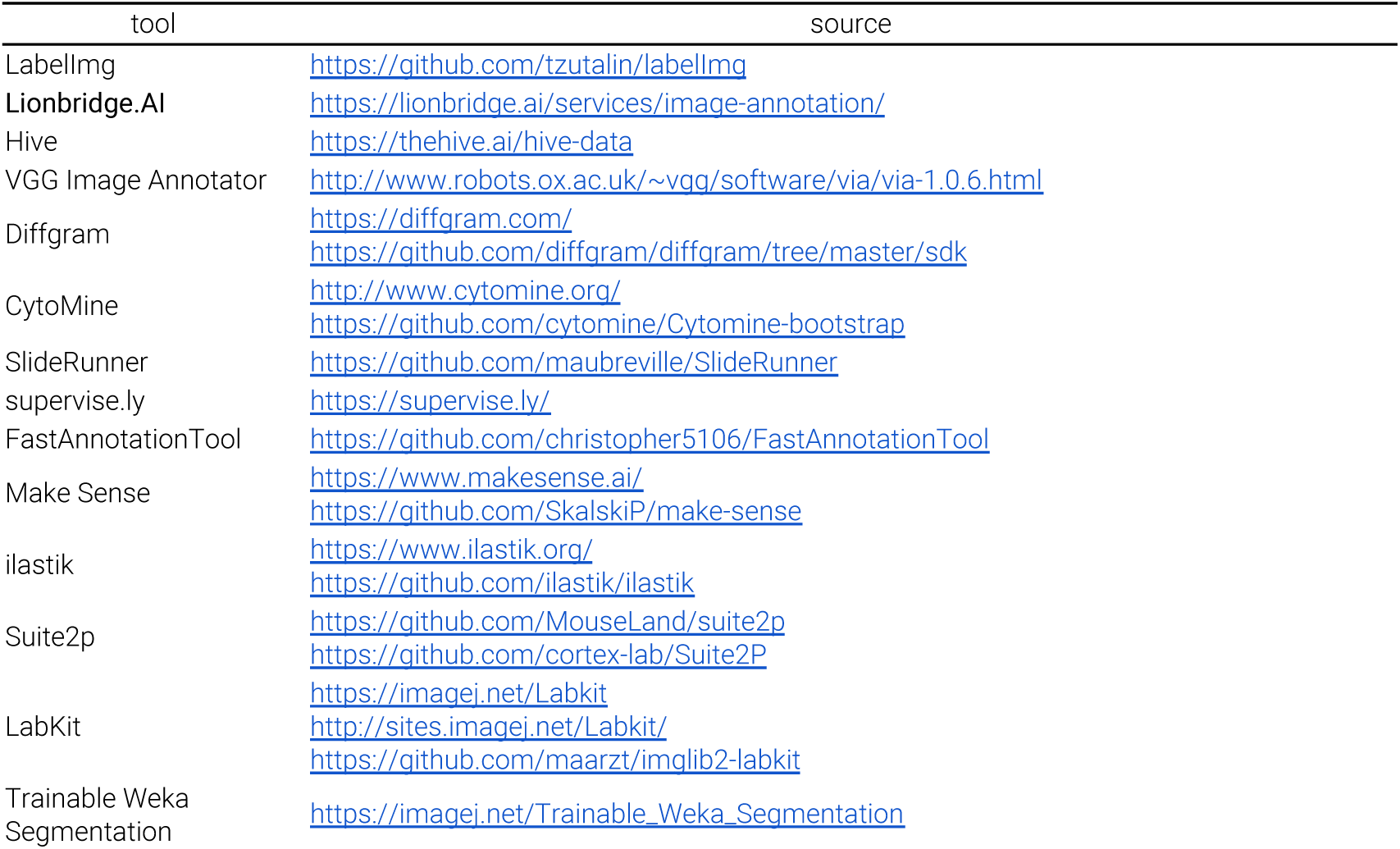
Sources of the compared software tools.

## Supplementary video

A short demonstration of *Contour assist* is provided as the video *figure2.mov*; available in higher resolution at https://drive.google.com/file/d/1dk3FrX-KIhTpaNYSKv-zVsoAoVUGtEYp/view?usp=sharing.

## Notes

### Competing Interest Statement

The authors have declared no competing interest.

### Summary of Updates

New experiments were performed on general image data (cars) and additional microscopy data (electron microscopy) that support the general applicability of the plugin. Additional features are also described. More tools are listed in the comparative analysis.

